# Targeting MDSC-HTR2B to Improve Immune Checkpoint Inhibitors in Breast to Brain Metastasis

**DOI:** 10.1101/2024.06.20.599939

**Authors:** Mukund Iyer, Diganta Das, Aaron G. Baugh, Priya Shah, Brooke Nakamura, Saman Sedighi, Max Reed, Julie Jang, Frances Chow, Evanthia Roussos Torres, Josh Neman

## Abstract

Myeloid Derived Suppressor Cells (MDSCs) support breast cancer growth via immune suppression and non-immunological mechanisms. Although 15% of patients with breast cancer will develop brain metastasis, there is scant understanding of MDSCs’ contribution within the breast-to-brain metastatic microenvironment. Utilizing co-culture models mimicking a tumor-neuron-immune microenvironment and patient tissue arrays, we identified serotonergic receptor, HTR2B, on MDSCs to upregulate pNF-κB and suppress T cell proliferation, resulting in enhanced tumor growth. *In vivo* murine models of metastatic and intracranial breast tumors treated with FDA-approved, anti-psychotic HTR2B antagonist, clozapine, combined with immunotherapy anti-PD-1 demonstrated a significant increase in survival and increased T cell infiltration. Collectively, these findings reveal a previously unknown role of MDSC-HTR2B in breast-to-brain metastasis, suggesting a novel and immediate therapeutic approach using neurological drugs to treat patients with metastatic breast cancer.

## Main Text

Brain metastases (BM), with an incidence ranging from 8.3 to 14.3 per 100,000 individuals, are the most prevalent type of intracranial neoplasm *(1, 2)*. Despite advancements in therapy, the median survival for patients with breast-to-brain metastases remains a bleak 3-19 months *(3)*. Historically, the brain has been considered an immunological sanctuary that limited entry of peripheral immune cells. However, recent evidence indicates that brain metastases actively recruit bone marrow-derived myeloid cells (BMDMs), which can constitute 30-50% of the brain tumor mass *(4, 5)*. Among these BMDMs are myeloid-derived suppressor cells (MDSCs), a diverse group of immature myeloid cells known for their immunosuppressive and tumor-promoting activities, such as enhancing neovascularization, stemness, invasion, proliferation, and colonization *(6-9)*.

The brain metastatic environment presents a distinct challenge because of the ability of central nervous system (CNS)-resident cells to act as local mediators, influencing the plasticity and differentiation of invading immune cells *(10)*. Notably, neurons play a role in cellular signaling through the secretion of classical neurotransmitters such as GABA, glutamate, acetylcholine, dopamine, and serotonin, with tumors exploiting these neurotransmitters to facilitate their adaptation within the CNS *(11)*. The role of serotonin in neuroimmune circuits regulating inflammation and immunity is an emerging area of research *(12-14)*. Furthermore, the serotonergic receptor HTR2B can activate transcription factor NF-κB in cardiomyocytes and lung vasculature *(15, 16)*. In metastatic tumors, NF-κB regulates several pro-tumorigenic processes, including cell growth and survival, inflammation, and immune response *(17)*. If this HTR2B-NF-κB signaling axis is activated in MDSCs residing within the brain metastatic microenvironment, it could have significant implications for both tumor progression and immunosuppression.

In this study, we utilized co-culture models mimicking a tumor-neuron-immune microenvironment along with patient tissue arrays to elucidate the functional role of HTR2B-mediated NF-κB activation in MDSCs in the BM microenvironment in driving inflammation, T cell suppression, and breast cancer cell growth and colonization. Treatment with FDA-approved HTR2B antagonist, clozapine, concurrently with immune checkpoint inhibitor, anti-PD-1, increased tumor-infiltrating T cells, increased survival, and decreased tumor burden in *in vivo* models of metastatic and intracranial breast tumors. Targeting the HTR2B-NF-κB axis in MDSCs presents a novel and timely therapeutic strategy for metastatic breast cancer.

### MDSCs are present in the breast-to-brain microenvironment and upregulate NF-κB signaling

While the presence and function of monocytic (M-MDSCs) and granulocytic MDSCs (G-MDSCs) has been studied in primary brain tumors, they have yet to be examined in BM *(18)*. To explore whether MDSCs are present in human BM, we stained patient brain metastatic tissue microarrays for M-MDSCs (CD11b+ CD14+ CD15-HLA-DR-) and G-MDSCs (CD11b+ CD14-CD15+ HLA-DR-). We found both M-MDSCs and G-MDSCs present in breast-(Fig. 1A) and lung-(fig. S1A) to-brain metastases. In parallel, we found infiltration of M-MDSCs (CD45+, CD11b+, Ly6c+, Ly6g-) and G-MDSCs (CD45+, CD11b+, Ly6c-, Ly6g+) in a syngeneic model of 4T1 brain metastases (Fig. 1B, fig. S1B).

**Figure 1:**
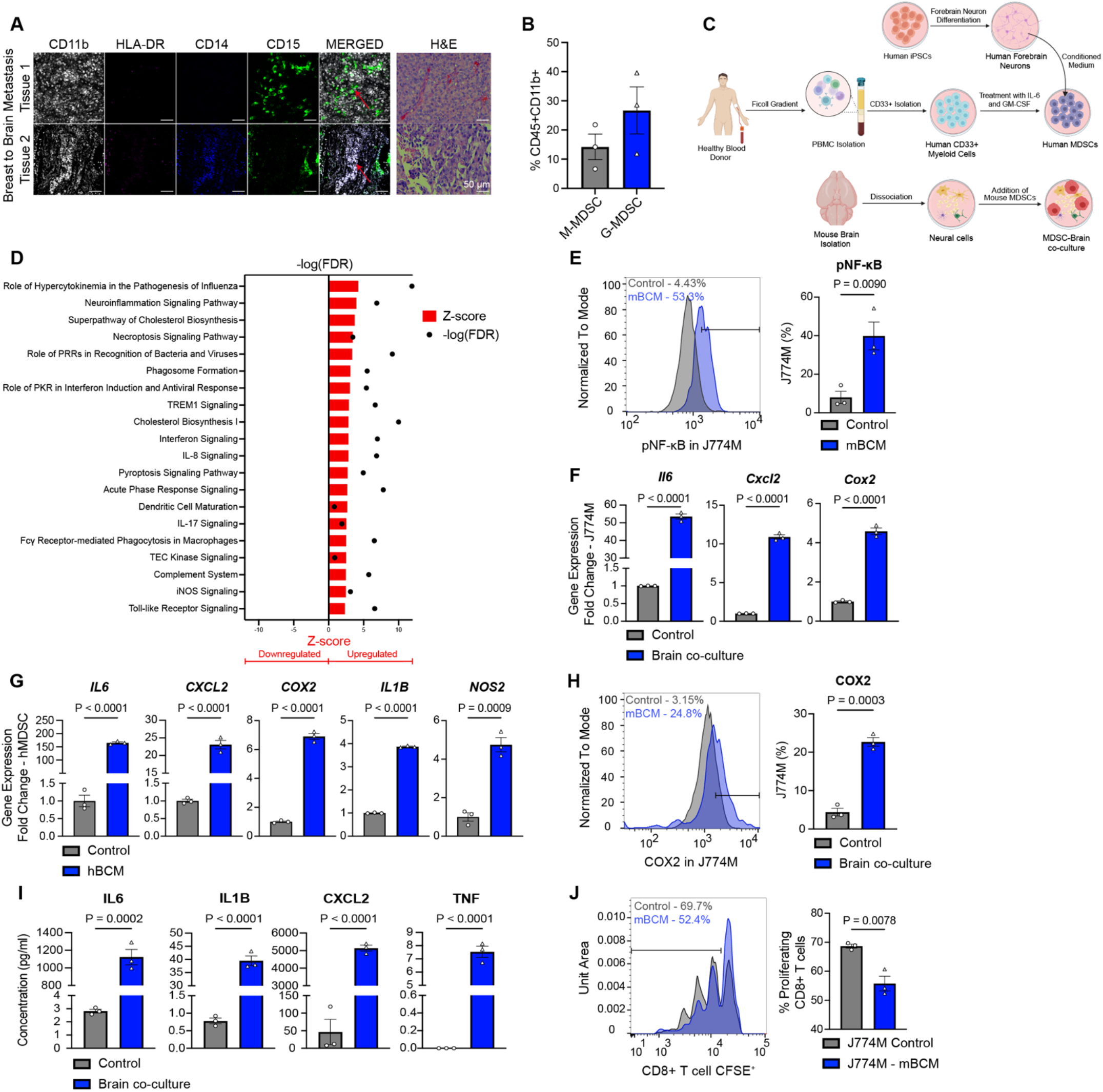
Brain metastatic MDSCs exhibit elevated pNF-κB signaling. **(A)** Immunofluorescence (IF) and H&E of breast-to-brain metastatic patient tissues. M-MDSCs (CD11b+ CD14+ CD15-HLA-DR-) and G-MDSCs (CD11b+ CD14-CD15+ HLA-DR-). Arrows point to MDSCs. Scale bar = 50 μm. **(B)** Flow cytometry of M- and G-MDSCs from 4T1 brain metastases 14 days post-transplantation. **(C)** Schematic representation of MDSC-brain microenvironment models. Top: Conditioned media was collected from induced pluripotent stem cell (iPSC)-derived human forebrain neurons and added to human MDSCs (hMDSCs) differentiated from peripheral blood mononuclear cells (PBMCs) of healthy donors (hMDSC-brain model). Bottom: Mouse MDSC J774M cells were co-cultured with neural cells isolated from postnatal mouse brains (mMDSC-brain model).**(D)** Ingenuity Pathway Analysis of significantly upregulated pathways in J774M in mMDSC-brain model by bulk RNA sequencing. **(E)** Flow cytometric analysis of pNF-κB expression in J774M treated with mouse brain conditioned media (mBCM). qPCR of NF-κB associated inflammatory markers in the mMDSC-**(F)** and hMDSC-**(G)** brain models. Flow cytometric analysis **(H)** and ELISA **(I)** of NF-κB associated inflammatory markers in the mMDSC-brain model. **(J)** CFSE proliferation assay of mouse CD8+ T cells co-cultured with J774M in control or mBCM conditions at a ratio of 4:1 (T cell:J774M). Each peak represents a cell division as determined by CFSE dilution. Percentage represents percent proliferating CD8+ T cells. Error bars represent ± SEM.

The immunologic landscape and tumor microenvironments between different organs distinctly influence MDSC plasticity and differentiation *(19)*. To understand MDSC adaptation and phenotypic plasticity within the brain microenvironment, we created murine and human *in vitro* models where we co-cultured mouse MDSC-like J774M cells with isolated mouse neural cells (mMDSC-brain model) and cultured human peripheral blood mononuclear cell-(PBMC-) derived MDSCs with human iPSC-derived forebrain neuron conditioned media (hMDSC-brain model; Fig. 1C, fig. S1C-D) *(20, 21)*. Differential gene expression analyses from bulkRNA-seq of isolated MDSCs from these neural-immune model revealed an upregulation of inflammatory and hypercytokinemia pathways when compared to MDSCs cultured alone (Fig. 1D).

Inflammatory cytokine expression is frequently driven by activation of the NF-κB transcription family, via nuclear translocation of cytoplasmic complexes *(22, 23)*. NF-κB activation in MDSCs enhances immunosuppressive capacity and promotes tumor survival and growth *(24, 25)*. We therefore hypothesized that MDSCs exhibit increased activation of NF-κB within the brain microenvironment. Indeed, MDSCs in our mMDSC- and hMDSC-brain models upregulated phospho-NF-κB (pNF-κB) and downstream inflammatory signals (Fig. 1E-I, fig. S1E-G). Furthermore, neural conditioned MDSCs exhibited an increased capacity to suppress CD8+ T cell proliferation (Fig. 1J), enhancing their immunosuppressive capabilities. These findings suggest a novel, pro-tumorigenic phenotype in MDSCs within the brain metastatic environment.

### Serotonin receptor, HTR2B, upregulates NF-κB signaling in MDSCs

Prior studies have demonstrated that the dynamic interplay between neuronal circuitry and myeloid cells in inflammation and extracranial cancers can alter cell plasticity and function *(26)*. Therefore, we hypothesized that paracrine neurotransmitters could influence pNF-κB upregulation in MDSCs within the brain microenvironment. Indeed, MDSCs treated with exogenous serotonin (5-HT) revealed a significant increase in pNF-κB expression relative to acetylcholine, dopamine, GABA, or norepinephrine (Fig. 2A, fig. S2A).

**Figure 2:**
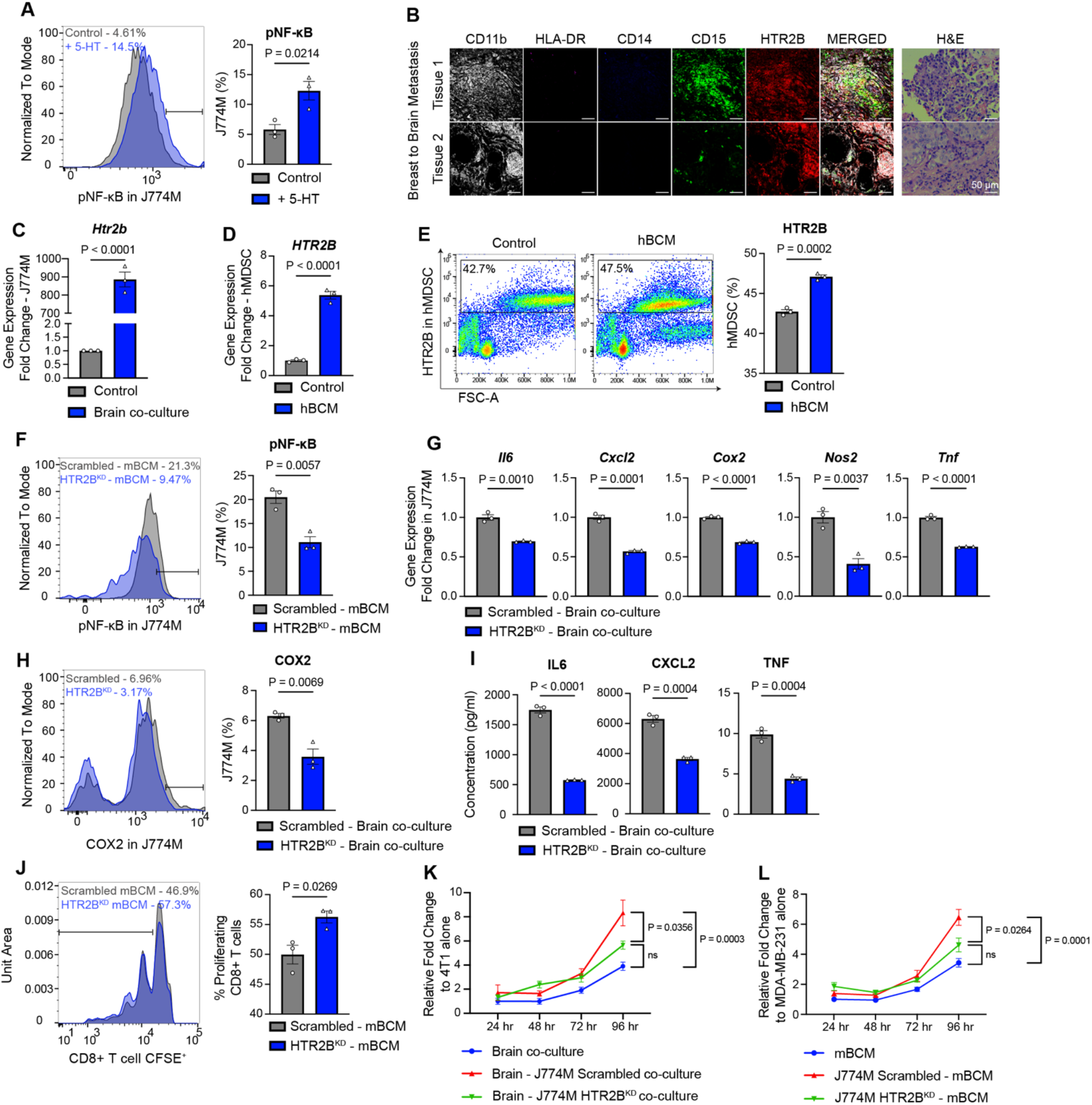
HTR2B regulates NF-κB signaling pathway in MDSCs. **(A)** Flow cytometric representative image and analysis of pNF-κB in J774M treated with 50 nM serotonin (5-HT) **(B)** IF and H&E staining of MDSCs and HTR2B in breast-to-brain metastatic patient tissues. Scale bar = 50 μm. qPCR analysis of HTR2B expression in the mMDSC-**(C)** and hMDSC-**(D)** brain models. **(E)** Flow cytometric analysis of HTR2B expression in the hMDSC-brain model. **(F)** Flow cytometric analysis of pNF-κB in J774M transduced with either shRNA control (Scrambled) or shRNA targeting HTR2B (HTR2B^KD^) after treatment with mBCM. **(G)** qPCR analysis of NF-κB associated inflammatory markers in J774M Scrambled or HTR2B^KD^ in the mMDSC-brain model. Flow cytometric analysis **(H)** and ELISA **(I)** of NF-κB associated inflammatory markers in J774M Scrambled or HTR2B^KD^ in the mMDSC-brain model. **(J)** CFSE proliferation assay of CD8+ T cells co-cultured with J774M Scrambled or HTR2B^KD^ at a ratio of 4:1 (T cell:J774M) after treatment with mBCM. **(K)** Proliferation assay of 4T1 breast cancer cells co-cultured with mouse neural cells and either J774M Scrambled or HTR2B^KD^. Data represent n=3-9 replicates per group per timepoint. **(L)** Proliferation assay of MDA-MB-231 breast cancer cells co-cultured with either J774M Scrambled or HTR2B^KD^ in mBCM. Data represent n=5-10 replicates per group per timepoint. Error bars represent ± SEM.

The immunomodulatory function of serotonin on MDSCs remains largely unexplored. HTR2B, a serotonergic receptor, has been shown to promote NF-κB activation in lung vasculature and cardiomyocytes (15, 16). Hence, we investigated serotonin receptor HTR2B expression in MDSCs within the brain microenvironment. We observed HTR2B expression in both M-MDSCs and G-MDSCs in human breast- and lung-to-brain metastatic patient tissues (Fig. 2B, fig. S2B). Furthermore, HTR2B expression was significantly upregulated in both our mMDSC and hMDSC-brain models (Fig. 2C-E, fig. S2C). These findings led us to hypothesize that MDSCs upregulate pNF-κB via an HTR2B-mediated mechanism. To test this hypothesis, we generated and validated shRNA-HTR2B knockdown (HTR2B^KD^) and shRNA-Control (Scrambled) mouse MDSC cells (fig. S2D-E). HTR2B^KD^ MDSCs in neural conditioned media displayed a significant downregulation of pNF-κB expression (Fig. 2F). Additionally, HTR2B^KD^ MDSCs in our mMDSC-brain model exhibited a significant downregulation of NF-κB downstream inflammatory markers (Fig. 2G-I, fig. S2F-H). These findings suggest that HTR2B regulates NF-κB activation and suppressive signaling within MDSCs residing in the brain microenvironment.

Given the known roles of NF-κB activation and its downstream effectors in regulating immune suppression, we investigated the impact of the HTR2B-NF-κB signaling axis on MDSC immunosuppressive function *(27, 28)*.Neural conditioned HTR2B^KD^ MDSCs demonstrated a decreased capacity to suppress CD8+ T cell proliferation, reducing their immunosuppressive capabilities (Fig. 2J). Beyond immunosuppression, MDSCs can enhance tumor cell proliferation *(29)*. Moreover, NF-κB downstream cytokines, IL-6 and TNF, have shown to stimulate cancer cell growth *(30, 31)*. Therefore, we hypothesized that MDSC-HTR2B signaling might influence breast tumor proliferation. In the brain microenvironment, 4T1 and MDA-MB-231 breast cancer cells co-cultured with HTR2B^KD^ MDSCs exhibited significantly decreased proliferative capacity (Fig. 2K-L).Collectively, these results support a critical mechanistic role for the HTR2B-NF-κB signaling axis in shaping both the immunological and non-immunological functions of MDSCs within the breast-to-brain metastatic microenvironment.

Furthermore, breast-to-brain metastases adapt to the CNS microenvironment through upregulation of neuronal characteristics for successful colonization and macro-metastatic growth *(11, 32)*. However, this examination of neuronal adaptation has overlooked the role of immune cells, primarily relying on neuron-tumor co-culture and immunodeficient mouse models. In the brain microenvironment, breast cancer 4T1 cells downregulate neuronal gene expression when co-cultured with MDSC Scrambled. However, this downregulation was mitigated when 4T1 were co-cultured with MDSC HTR2B^KD^ (fig. S3A). Specifically, we observed a significant decrease in nuclear colocalization of SRRM4 and expression of ABAT and Reelin, which are key neuronal signatures previously shown to be involved in the neural acquisition of tumors, in 4T1 co-cultured with MDSC Scrambled as compared to MDSC HTR2B^KD^ (fig. S3B) (11, 32). These findings suggest MDSC-HTR2B plays a critical role in regulating neural acquisition and colonization of breast tumor cells seeking to exploit the CNS microenvironment.

### HTR2B antagonism delays brain metastasis incidence leading to increased survival

To assess the potential clinical applicability of these findings, we investigated targeting the HTR2B-NF-κB signaling pathway using small molecule inhibitors. There is a diverse array of HTR2B antagonists available, including FDA-approved drugs that are capable of crossing the blood-brain barrier *(33)*. Both human and mouse MDSCs exposed to neural conditioned media downregulated NF-κB downstream inflammatory signals when treated with HTR2B antagonists. (Fig. 3A, fig. S4A-B). These results suggest that HTR2B antagonists can effectively mitigate the HTR2B-NF-κB signaling axis in MDSCs, leading to a reduction in pro-tumorigenic markers. Furthermore, our results demonstrate that MDSC-HTR2B knockdown within the brain microenvironment reduces tumor growth and promotes T cell proliferation, creating a potentially favorable environment for enhanced anti-tumor immunity. Therefore, we hypothesized that combining FDA-approved HTR2B antagonist, clozapine, with immune checkpoint inhibitor, anti-PD-1, could synergistically augment the anti-tumor response leading to increased survival and reduced metastatic brain tumor burden. To test this hypothesis, syngeneic metastatic breast tumor mice were concurrently treated with clozapine and anti-PD-1 (Fig. 3B). Mice receiving this dual treatment exhibited a significant increase in survival compared to those receiving either single-agent or vehicle treatment (Fig. 3C). Specifically, the median survival of vehicle-treated mice was 16 days post-tumor seeding, whereas mice treated with both clozapine and anti-PD-1 survived for 25 days – a 56% increase. Additionally, there was a reduction in overall tumor burden in mice receiving either anti-PD-1 treatment alone or clozapine + anti-PD-1 treatment compared to vehicle treatment or clozapine treatment alone (Fig. 3D, fig. S4C). Specifically, the dual treatment group displayed a significantly delayed median onset (Fig. 3E) and decreased burden (Fig. 3F) of brain metastases compared to vehicle treatment or clozapine treatment alone. While anti-PD-1 treatment alone yielded a reduction in overall tumor burden comparable to the dual treatment with clozapine and anti-PD-1, it did not significantly affect brain tumor burden. All together, these findings suggest a strong synergistic effect of HTR2B antagonism with anti-PD-1 specifically in reducing brain metastasis.

**Figure 3:**
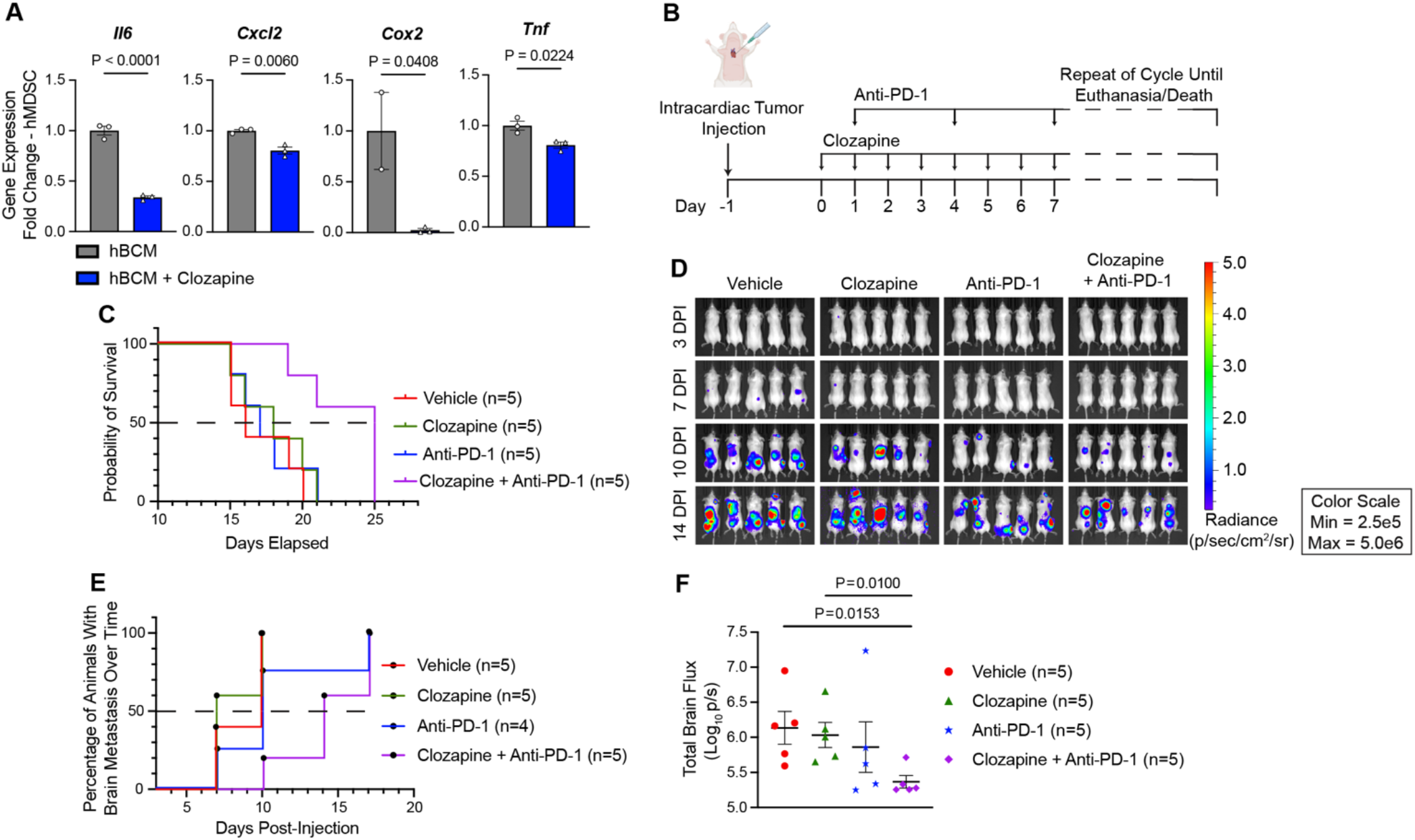
HTR2B antagonism and immunotherapy synergistically delay breast-to-brain metastasis and improve survival. **(A)** qPCR analysis of NF-κB associated inflammatory markers in hMDSCs treated with 10 uM clozapine in hBCM. **(B)** *In vivo* mouse breast metastatic model. Female mice were injected intracardially with 4T1 breast cancer cells and subsequently treated with either vehicle control, clozapine (administered daily), anti-PD-1 (administered twice per week), or a combination of clozapine and anti-PD-1. **(C)** Kaplan-Meier survival curve of mice described in (B). Mice in the clozapine + anti-PD-1 group exhibited improved survival compared to vehicle (p=0.0086, hazard ratio [HR]=11.25), clozapine alone (p=0.0244, HR=6.715), and anti-PD-1 alone (p=0.0136, HR=8.601) (log-rank test). n=5 per group. **(D)** *In vivo* bioluminescence imaging (BLI) of mice described in (B) at 3,7,10, and 14 days post-intracardiac tumor cell injection. BLI intensity scale was normalized (Min: 2.5e^5^; Max:5.0e^6^). **(E)** Graphical comparison of brain metastasis (BM) incidence over time in animals (n=5 per group). BM incidence determined through BLI signal. Mice in the clozapine + anti-PD-1 group displayed a delayed onset of BM compared to vehicle (p=0.0116, HR=12.81) and clozapine alone (p=0.0080, HR=14.38) (log-rank test). **(F)** Bar graph represents mouse head BLI signal at day 10 post-tumor implantation. Error bars represent ± SEM.

### HTR2B inhibition increases survival via enhanced T cell infiltration within intracranial tumor

Next, we aimed to understand whether the observed synergy between clozapine and anti-PD-1 treatment results in increased T cell infiltration in the brain microenvironment. To investigate this, mice intracranially implanted with breast cancer were treated concurrently with clozapine and anti-PD-1. (Fig. 4A). We observed a significant increase in survival only in mice treated with clozapine and anti-PD-1 but not with either single-agent treatment (Fig. 4B). Median survival in vehicle-treated mice were 20 days post-tumor seeding, while mice receiving both clozapine and anti-PD-1 treatment exhibited a 30% increase, surviving for 26 days. The metastatic brain tumors of these mice revealed a significant increase in CD3+ T cell infiltration in those treated with both clozapine and anti-PD-1 compared to vehicle or single agent treatment groups (Fig. 4C). These findings suggest that combining HTR2B antagonism with immunotherapy could transform the breast-to-brain tumor microenvironment from immunologically “cold” to “hot,” potentially leading to improved survival outcomes.

**Figure 4:**
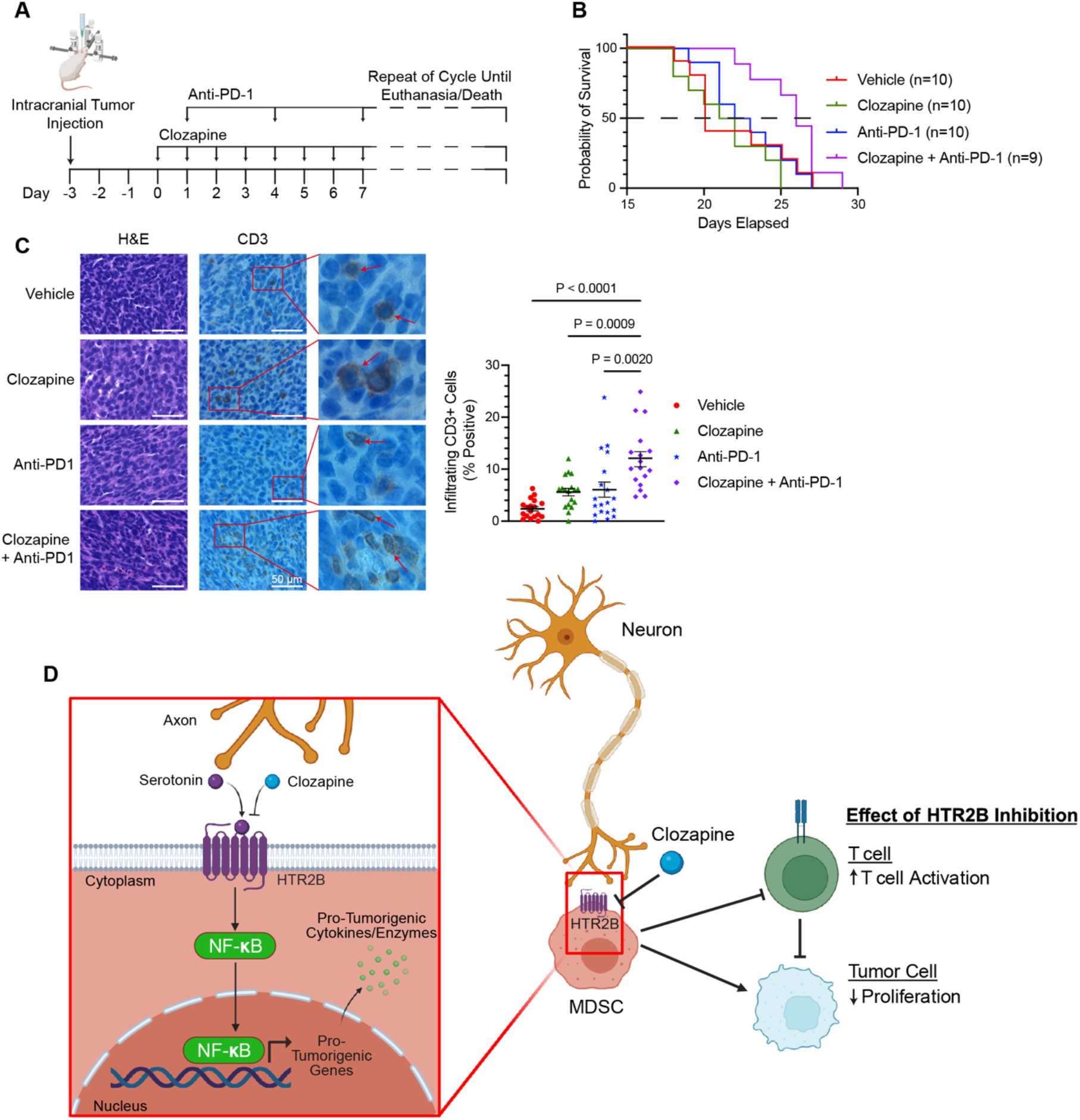
HTR2B inhibition and immunotherapy promotes T cell infiltration into intracranial tumor. **(A)** Mouse breast-to-brain metastatic model. Female mice were injected intracranially with 4T1 breast cancer cells and subsequently treated with either vehicle control, clozapine (administered daily), anti-PD-1 (administered twice per week), or a combination of clozapine and anti-PD-1. **(B)** Kaplan-Meier survival curve of mice described in (A). Mice in the clozapine + anti-PD-1 group demonstrated improved survival compared to vehicle (p=0.019, HR=4.001), clozapine alone (p=0.0009, HR=7.849), and anti-PD-1 alone (p=0.0193, HR=3.727) (log-rank test). n=9-10 per group. **(C)** IHC images and analysis of brain tumor tissue sections stained for CD3+ T cells across the groups outlined in (A). H&E staining is included. Arrows indicate CD3+ cells. 3 mice per group, 5-6 images per tissue. Scale bar = 50 μm. **(D)** Schematic illustration of the HTR2B-NF-κB signaling axis in MDSCs within the metastatic brain microenvironment. Error bars represent ± SEM.

Our findings collectively suggest a novel mechanism by which MDSCs in the brain microenvironment leverage extracellular serotonin to activate a pro-tumorigenic signaling cascade through HTR2B. This HTR2B-mediated activation of NF-κB suppressive signaling leads to enhanced T cell suppression and promotion of tumor proliferation. However, HTR2B antagonism with clozapine disrupts this suppressive pathway, resulting in restoration of T cell activation and reduction of tumor proliferation (Fig. 4D).

## Discussion

This study contributes to the emerging field of cancer neuroscience by elucidating an intricate communication between tumor, nervous, and immune systems in brain metastasis. Prior research has primarily focused on how the brain influences tumor cells *(13, 34)*. Our work reveals how MDSCs adapt to and exploit the unique brain signaling environment to exert pro-tumorigenic effects. In response to neural cues, MDSCs exhibit upregulated NF-κB signaling, driven by paracrine serotonin through the serotonergic receptor HTR2B. The HTR2B-NF-κB signaling axis is crucial for MDSC-induced immunosuppression, tumor cell proliferation, and the integration of tumors into CNS. Targeting this axis with the FDA-approved HTR2B antagonist clozapine, in combination with the immune checkpoint inhibitor anti-PD-1, results in a significant reduction in overall tumor burden, increased survival, and decreased brain metastasis through enhanced T cell infiltration.

Our findings confirm the established role of MDSC-mediated immunosuppression and the pro-tumorigenic function of NF-κB in cancer *(7-10, 19)*. Distinctively, our study identifies HTR2B as a key mediator of NF-κB activation in MDSCs. Interestingly, while increased tumor proliferation was observed with MDSC-HTR2B activity, there was a concurrent decrease in tumor assimilation within the brain microenvironment. This is particularly notable because tumor cells are known to upregulate neuronal genes to survive within the brain *(11, 32)*. Therefore, MDSC-HTR2B signaling in the brain microenvironment can create a pro-tumorigenic niche, allowing tumor cells to thrive without requiring drastic phenotype changes to mimic neural cells.

Previous research utilizing immune checkpoint inhibitors against breast-to-brain metastasis has shown limited efficacy *(34)*. In contrast, our work demonstrates that priming the immune microenvironment with the FDA-approved HTR2B antagonist, clozapine, combined with concurrent administration of the anti-PD-1 checkpoint inhibitor, leads to a significantly more effective anti-tumor response. This approach, leveraging an existing FDA-approved medication, underscores the strong potential for our findings to be translated into clinical applications, benefiting both patients with metastatic breast cancer and those with established brain metastases.

In conclusion, our study reveals a novel HTR2B-NF-κB signaling axis in MDSCs that promotes breast-to-brain metastasis. Targeting this axis with HTR2B antagonists holds promise for developing novel therapeutic strategies to enhance immunotherapy efficacy and address the challenges of treating brain metastases.Additionally, these findings prompt a need to rethink the use of neurological drugs to further promote and optimize immunotherapy treatments.

## Acknowledgments

We would like to acknowledge patient advocates and all breast cancer patients/survivors/family members for the invaluable role they have played in our research over the years. We would also like to thank Ivetta Vorobyova and Bernadette Masinsin for help with core-services at USC. The publication was supported in part by award number P30CA014089 from the National Cancer Institute. The content is solely the responsibility of the authors and does not necessarily represent the official views of the National Cancer Institute or the National Institutes of Health.

## Funding

National Institutes of Health/National Cancer Institute R01CA223544-01A1 (JN)

Department of Defense BCRP BC141728 (JN)

National Institutes of Health T90 DE021982 (MI)

National Center for Advancing Translational Science KL2TR001854 (FC)

American Cancer Society IRG-22-144-60 (FC)

## Author contributions

Conceptualization: MI, DD, JJ, ERT, JN

Methodology: MI, DD, BN, AGB, JJ, FC, ERT, JN

Investigation: MI, DD, MR, PS, AGB, BN, SS, FC

Visualization: MI, PS, MR

Funding acquisition: MI, FC, ERT, JN

Project administration: ERT, JN

Supervision: JN

Writing – original draft: MI, JN

Writing – review & editing: MI, DD, AGB, PS, BN, SS, MR, JJ, FC, ERT, JN

## Data and materials availability

All data are available in the main text or the supplementary materials.

## Supplementary Materials

Materials and Methods

Figs. S1 to S4

Tables S1 to S2

