## supplemental file for "Targeting MDSC-HTR2B to Improve Immune Checkpoint Inhibitors in Breast to Brain Metastasis"

### Materials and Methods

#### Lentiviral Particle Production, Transduction, and Selection

*HTR2B* knockdown and Scrambled (GeneCopoeia, Cat# MSH027376-LVRU6P) and pHIV-Luc-ZsGreen (Addgene, plasmid no. 39196) lentiviral particles were produced using the 293T cell system as previously described (35). J774M cells were transduced to stably express knockdown ($HTR2B^{KD}$) or Scrambled. These cells were treated with 4 µg/ml puromycin to ensure proper cell selection. Three J774M $HTR2B^{KD}$ variants were tested and one variant with significant knockdown was chosen to be utilized in future experiments. 4T1 and MDA-MB-231 cells were transduced to stably express Firefly Luciferase/GFP (FF/GFP). These cells were FACS sorted for the GFP+ population.

#### Immunofluorescence

Brain-metastatic tumor tissues from patients were acquired via USC Neurosurgery under approved consent and I. These tissues were formalin fixed, paraffin embedded, and processed into 10 uM-thick sections. Selected regions of interest were punch-holed and embedded into a tissue microarray with 50+ samples. Immunofluorescence protocol on tissue and cells were performed as previously described (11). Primary antibodies and secondary antibodies list can be found in Supplementary Table 2.

#### Immunohistochemistry

Mouse brain tissues were fixed, paraffin embedded, and processed as described above. Tissues were stained on the Leica Bond III Autostainer (Leica, Cat# 21.2201). Antigen unmasking was conducted with BOND Epitope Retrieval Solution 1 & 2 (Leica, Cat# AR9961/AR9640) for 30 minutes as per manufacturer's instructions. Primary antibodies were incubated for 60 minutes. Subsequent use of a DAB chromogen and hematoxylin & eosin stain were conducted utilizing the BOND Polymer Refine Detection DAB (Leica, Cat# DS9800) and Modified Mayer’s Hematoxylin (American Mastertech, Cat# HXMMHGAL) as per manufacturer’s instructions.

#### Microscopy and Imaging

Confocal and widefield imaging were performed and quantified as previously described (32).

#### ELISA

In this study, we used Luminex xMAP technology for multiplexed quantification of 45 Mouse cytokines, chemokines and growth factors. The multiplexing analysis was performed using the Luminex™ 200 system (Luminex, Austin, TX, USA) by Eve Technologies Corp. (Calgary, Alberta). Forty-five markers were simultaneously measured in the samples using Eve Technologies' Mouse Cytokine 45-Plex Discovery Assay® which consists of two separate kits: one 32-plex and one 13-plex (MilliporeSigma, Burlington, Massachusetts, USA). The assay was run according to the manufacturer's protocol. The 32-plex consisted of Eotaxin, G-CSF, GM-CSF, IFNγ, IL-1α, IL-1β, IL-2, IL-3, IL-4, IL-5, IL-6, IL-7, IL-9, IL-10, IL-12(p40), IL-12(p70), IL-13, IL-15, IL-17, IP-10, KC, LIF, LIX, MCP-1, M-CSF, MIG, MIP-1α, MIP-1β, MIP-2, RANTES, TNFα, and VEGF. The 13-plex consisted of 6Ckine/Exodus2, Erythropoietin, Fractalkine, IFNβ-1, IL-11, IL-16, IL-20, MCP-5, MDC, MIP-3α, MIP-3β, TARC, and TIMP-1. Assay sensitivities of these markers range from 0.3 – 30.6 pg/mL for the 45-plex. A cubic spline and 5-paramter logistic regression were used when formatting and optimizing the standard. Regression analysis was performed utilizing the Bio-Plex Manager™ software. Individual analyte sensitivity values are available in the MilliporeSigma MILLIPLEX® MAP protocol.

#### Primary Mouse Brain Cells

Mouse brain cells were isolated from whole brain tissue of 0 - 4 day postnatal mice and cultured *in vitro* as previously described (36). Media collected from these cultures every 72 hours serve as mouse brain conditioned media (mBCM) for subsequent experiments.

**Human iPSC-derived Forebrain Neurons**

Human PGP-1 induced pluripotent stem cells (iPSCs) were kindly provided by Dr. Giorgia Quadrato. Neural progenitor cells (NPCs) were generated from PGP-1s utilizing the STEMdiff$^{TM}$ SMADi Neural Induction Kit (STEMCELL Technologies, cat# 08581) as per manufacturers instructions. Mature forebrain neurons were generated from NPCs utilizing the STEMdiff$^{TM}$ Forebrain Neuron Differentiation Kit (STEMCELL Technologies, cat# 08600) and STEMdiff$^{TM}$ Forebrain Neuron Maturation Kit (STEMCELL Technologies, cat# 08605) as per manufacturer's instructions. Conditioned media was collected from mature forebrain neurons between 8-20 days after incubation in neuron maturation medium.

**Tumor/MDSC-Neuron Co-cultures**

Tumor cells and MDSCs were stained with cell permeable Far Red dye (Thermo Fisher, catalog no. C34564) at 10 uM or CMFDA dye (Thermo Fisher, catalog no. C2925) at 5 uM as per manufacturers recommendation to allow for future separation of cell populations. Cells were resuspended in complete Neurobasal-A media and seeded onto neuronal cultures (1:1:125 tumor to MDSC to neuron ratio) to model Tumor-MDSC-Neuron interactions. 72 hours post seeding, half of the media was replaced. 96 hours post seeding, tumor cells/MDSCs were either separated via FACS for qPCR, stained and processed for flow cytometry, or fixed with 4% formaldehyde for immunofluorescence studies.

**Cell Culture**

The following commercially available cell lines were used: 4T1 triple negative breast cancer cells (ATCC), MDA-MB-231 triple negative breast cancer cells (ATCC). MDSC-like J774Ms were kindly provided from Dr. Kebin Liu's lab at Augusta University. All cell cultures were maintained at 37°C, 5% CO2 in humidified incubators. These commercial cell lines, differentiated Human MDSCs, and T cells were cultured in RPMI1640 (Thermo Fisher, catalog no. 11875119) supplemented with 10% FBS (Omega Scientific, catalog no. FB-12), 1.5% HEPES buffer (Gibco, catalog no. 25-060-Cl), 1x GlutaMAX (Thermo Fisher, catalog no. 35050061), 1% MEM nonessential amino acids (Corning, catalog no. 25-025-Cl), 1x Antibiotic-Antimycotic (Thermo Fisher, catalog no. 15240062), 1% sodium pyruvate (Gibco, catalog no. 11360-070), 0.0004% beta-mercaptoethanol (Sigma, catalog no. M3148) and will be referred to as complete RPMI media. Primary mouse neurons were maintained in Neurobasal-A Media (Thermo Fisher, catalog no. 10888022) supplemented with 1x B-27 (Thermo Fisher, catalog no. 17504044), 1x Antibiotic-Antimycotic (Thermo Fisher, catalog no. 15240062), and 1x GlutaMAX (Thermo Fisher, catalog no. 35050061) and will be referred to as complete Neurobasal-A media. All cell lines were stored in liquid nitrogen and frozen down between 5 to 10 passages from original cell line. All cells thawed from liquid nitrogen were passaged once before experiment setup and no cell lines were utilized past 15 passages. Cell lines were negative for mycoplasma and were frequently tested utilizing MycoAlert Kits (VWR, catalog no. 75860-360).

**Tumor model, treatment dosing scheme, and tumor resection**

To model metastatic disease, $5 \times 10^3$ triple-negative breast cancer 4T1 FF/GFP resuspended in serum-free RPMI1640 were injected intra-cardiac into the heart's left ventricle with a 25-gauge needle. Clozapine was dosed daily by intraperitoneal injection at 1 mg/kg body weight. Clozapine was diluted in 0.1N HCl and neutralized with 4.0N NaOH and diluted in PBS. Anti-PD-1 (BioXCell; RMP1-14) was diluted in PBS and dosed 2x/week by intraperitoneal injection at 100 ug/mouse. Development of brain metastasis and other distant metastasis was measured by optical imaging of animals twice per week. Luciferin (1μL/gram bodyweight of animal) was injected into the peritoneal cavity of each mouse and bioluminescence was imaged (dorsal and ventral) 10 minutes after injection. We measured bioluminescent signal from brain metastases utilizing the same brain ROI for all experimental animals on all imaging days. To model brain metastatic disease, $5 \times 10^3$ 4T1 resuspended in serum-free RPMI1640 were injected into the brain parenchyma utilizing a stereotaxic frame. After locating the intersection point of bregma and midline (x = 0, y = 0, z = 0), tumor cells were injected with a 25-guage needle at (x = +1.0, y = +1.0, z = -1.0). Clozapine and anti-PD-1 were dosed as described above. Tumors were seeded for 3 days prior to beginning treatment. Mice for all experiments were monitored and weighed daily for presentation of tumor burden related symptoms and were humanely euthanized.

Immediately post-euthanasia, whole brains were resected and placed into formalin before being sent to the histology core at USC for paraffin embedding and processed into 10 uM-thick sections for subsequent staining.

**Animals**

9-12 week-old adult female BALB/c mice (strain#000651) were purchased from Jackson Laboratories and used for either *in vivo* intracardiac or intracranial injection experiments to model metastatic disease. Animal procedures were performed under approved IACUC protocols and guidelines. All animals were humanely euthanized upon signs of morbidity, including development of tumor symptoms (paralysis, hydrocephalus, weight loss, severely hunched).

**In vitro suppression assay**

CD8+ T cells were isolated from spleens of healthy 8-12 week female BALB/c mice via the EasySep Mouse CD8+ Isolation Kit (Stem Cell, catalog no. 19853) per manufacturer's instruction. Isolated CD8+ T cells were stained with CFSE at 3 uM. $5x10^4$ CFSE-labeled $CD8^+$ T cells were co-cultured with J774M cells (Control, Scrambled, $HTR2B^{KD}$) at a 4:1 ratio (T cell:J774M) and anti-CD3/CD28 beads (Thermo Fisher Scientific, catalog no. 11453D) as per manufacturer's instructions in complete RPMI media. Media was changed after 24 hours to mBCM. T cells were allowed to proliferate for a total of 60 hours. Afterwards, the samples were collected, stained with Live/Dead Fixable Aqua (Thermo Fisher, catalog no. L34965) and analyzed via flow cytometric analysis.

**Flow Cytometry**

To detect intracellular markers, The Foxp3/Transcription Factor Staining Buffer Set (Invitrogen, catalog no. 00-5523-00) was utilized to fix and permeabilize cells as per manufacturers recommendation. Cells were subsequently blocked in CD16/CD32 Fc block (BD Biosciences; catalog no.553142) at a 1:100 dilution overnight. The next day, samples were incubated with primary antibody and secondary antibodies (Supplementary Table 2) and ran on the Attune NxT Flow Cytometer at the USC Stem Cell Flow Cytometry Facility. To detect extracellular markers, cells were not fixed or permeabilized but rather directly stained with primary and secondary antibodies and resuspended in FACS buffer (5% FBS, 0.1% NaN3, PBS) prior to running on the Attune NxT. For Fluorescent Activated Cell Sorting (FACS), previously stained CMFDA or Far-Red cells were harvested, resuspended in FACS buffer, and analyzed on the BD FACSAria Cell Sorter. Resulting data was analyzed utilizing FlowJo v10.10 software.

**RNA Isolation and qPCR Analysis**

Cells were harvested by scraping when appropriate or trypsinization for 3-5 minutes at 37°C, followed by neutralization in media containing 10% FBS, and centrifugation at 360 rcf for 5 minutes. The resulting cell pellet was either processed for RNA extraction immediately or frozen for subsequent use in qPCR (preformed in triplicate per sample) as previously described (11). All primers utilized for qPCR analysis were purchased from IDT (Supplementary Table 1).

**Exogenous Neurotransmitter or HTR2B Antagonist Treatment**

J774M were seeded at $7.5 \times 10^4$ cells onto a 6 well plate in complete RPMI media. For exogenous neurotransmitter assays, the following day the media was replaced with complete Neurobasal-A Media and concurrently treated with Serotonin (Sigma-Aldrich, catalog no. H9523), Dopamine (Sigma-Aldrich, catalog no. H8502), Acetylcholine (Sigma-Aldrich, catalog no. A6625), Norepinephrine (Sigma-Aldrich, catalog no. A7256), or GABA (Sigma-Aldrich, catalog no. A2129). For HTR2B antagonist treatment assays, the following day the media was replaced with mouse BCM and concurrently treated with SB-204741 (Cayman Chemical, catalog no. 32965), clozapine (TCI Chemicals, catalog no. 2547), RS-127445 (Adooq Bioscience, catalog no. A11165) or Aripiprazole (TCI Chemicals, catalog no. A2496). Samples were collected for downstream RNA isolation/qPCR, flow cytometry, or ELISA.

**Proliferation Assay**

4T1 FF/GFP and J774M (Scrambled, HTR2B$^{KD}$) cells were resuspended in complete Neurobasal-A media and seeded onto neuronal cultures (1:1:125 tumor to MDSC to neuron ratio) on a 96 well plate. MDA-MB-231 FF/GFP and J774M (Scrambled, HTR2B$^{KD}$) cells were resuspended in complete RPMI media and seeded (2:1 tumor to MDSC ratio) on a 96 well plate. Media was changed the following day to mouse brain conditioned media (mBCM). Fluorescent GFP signal is indicative of the number of tumor cells in the well. Fluorescence was measured daily on a Varioskan™ Lux multimode microplate reader (ThermoFisher Scientific) at 485/520 (Ex/Em).

**RNA-seq of J774M cells**

J774M cells were cultured independently or in a MDSC-Neuron co-culture as described above and J774M were separated via FACS in singlet. Transcriptomic sequencing was performed at the USC Norris Molecular Genomics Core. Samples were simultaneously library prepped using the Kapa mRNA HyperPrep kit following manufacturer's protocol (Roche, KK8580). Prepared libraries were sequenced on the Illumina Nextseq500 at single end 75 cycles. RNA integrity number values of the samples ranged from 8.5 to 10. Samples were read at 25 million reads per sample at a read length of 1x75. Processing of the data was conducted on Partek Flow via DESeq2 to generate a differentially expressed gene list amongst samples. These differentially expressed genes were inputted into Ingenuity Pathway Analysis and analyzed for significantly enriched canonical signaling pathways.

**Human MDSC Derivation**

Blood from healthy donors was obtained at the University of Southern California under approved consent and IRB. Peripheral blood mononuclear cells (PBMCs) were separated via a Ficoll gradient and isolated for CD33+ cells via the EasySep Human CD33+ Selection Kit II (Stem Cell, catalog no. 17876) per manufacturer's instructions. These cells were plated at $5 \times 10^5$ cells/mL in complete RPMI media for 7 days in the presence of recombinant human GM-CSF (20 ng/mL; Stem Cell, catalog no. 78140) and IL-6 (20 ng/mL; Stem Cell, catalog no. 78050.1). GM-CSF was added on days 1, 3, and 5 whereas IL-6 was only added on day 5. After 7 days, the cells were utilized for downstream experiments and T cell suppression assays were preformed to validate the suppressive capability of these Human MDSCs.

**Statistical Analysis**

Statistics were preformed using GraphPad Prism 8 software. To assess statistical significance, Student's t test, one-way ANOVA, and log-rank statistical analyses were utilized. For one-way ANOVA, post hoc analysis was performed using Tukey's multiple comparison test where necessary. All histogram data show individual data points with the mean ± SEM (Standard Error of the Mean) and $p < 0.05$ was considered to be statistically significant. Statistical significance and Hazard Ratio for survival data from *in vivo* experiments were calculated using Log-Rank Test.

### Supplemental Figure 1

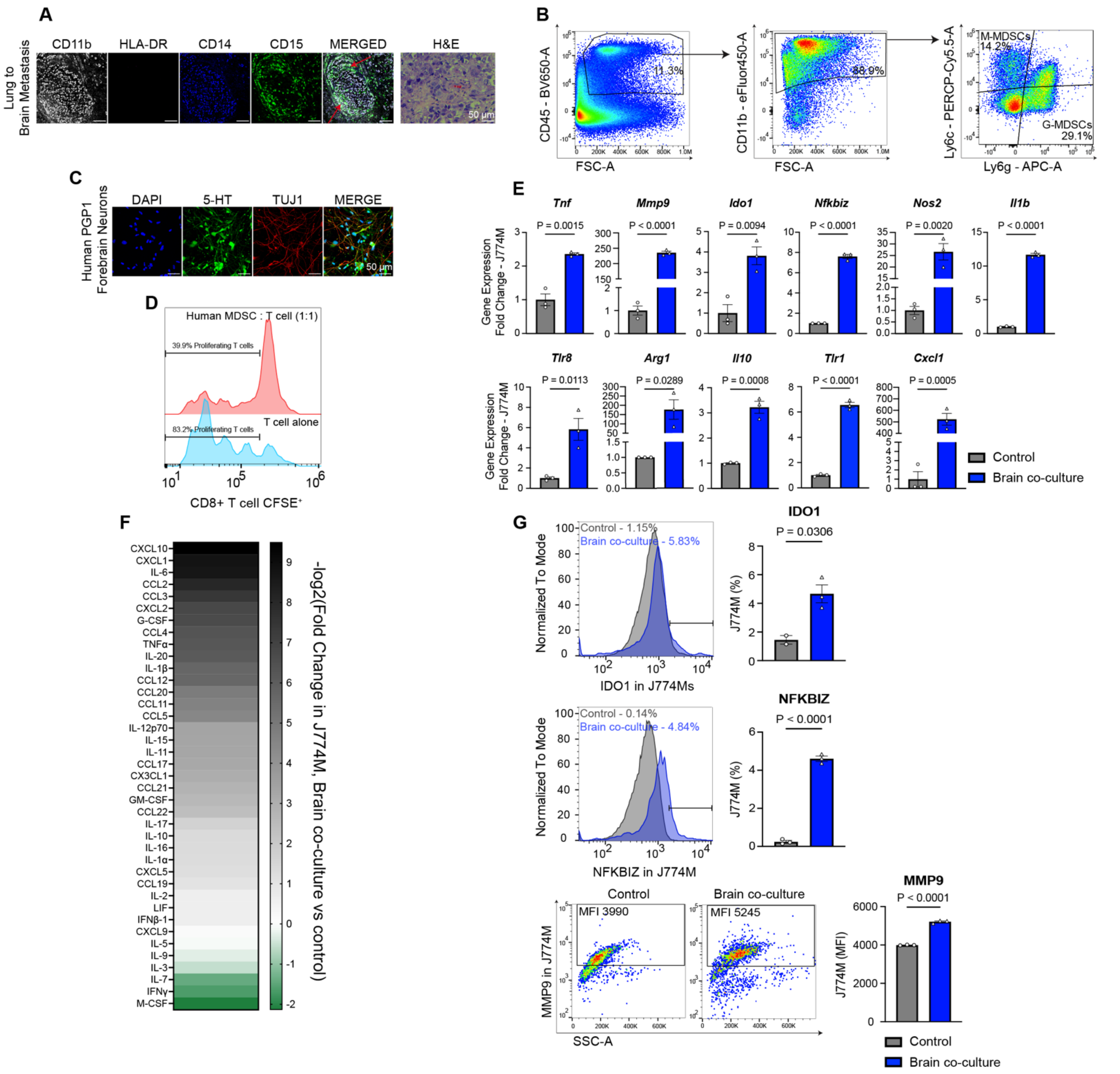

**Supplemental Figure 1: Elevated pNF-κB signaling in brain metastatic MDSCs. (A)** IF and H&E staining of MDSC markers in lung-to-brain metastatic patient tissues. Arrows point to MDSCs. Scale bar = 50 μm. **(B)** Flow cytometric gating strategy used to identify M-MDSCs and G-MDSCs in resected brain tumors from mice 14 days after intracranial injection of 4T1 breast cancer cells. **(C)** IF staining of human mature forebrain neurons differentiated from iPSCs generated using the PGP-1 cell line. Images captured 8 days post incubation in neuron maturation media. Scale bar = 50 μm. **(D)** CFSE proliferation assay of human CD3+ T cells co-cultured with hMDSCs at a 1:1 ratio. **(E)** qPCR analysis of NF-κB associated inflammatory markers and immunosuppressive markers in the mMDSC-brain model. **(F)** Heatmap depicting the results of an ELISA analysis comparing the levels of NF-κB associated inflammatory markers in the mMDSC-brain model. **(G)** Flow cytometric images and analysis of NF-κB associated inflammatory markers in the mMDSC-brain model. Signal measured either by percent positive or median fluorescent intensity (MFI). Error bars represent ± SEM.

### Supplemental Figure 2

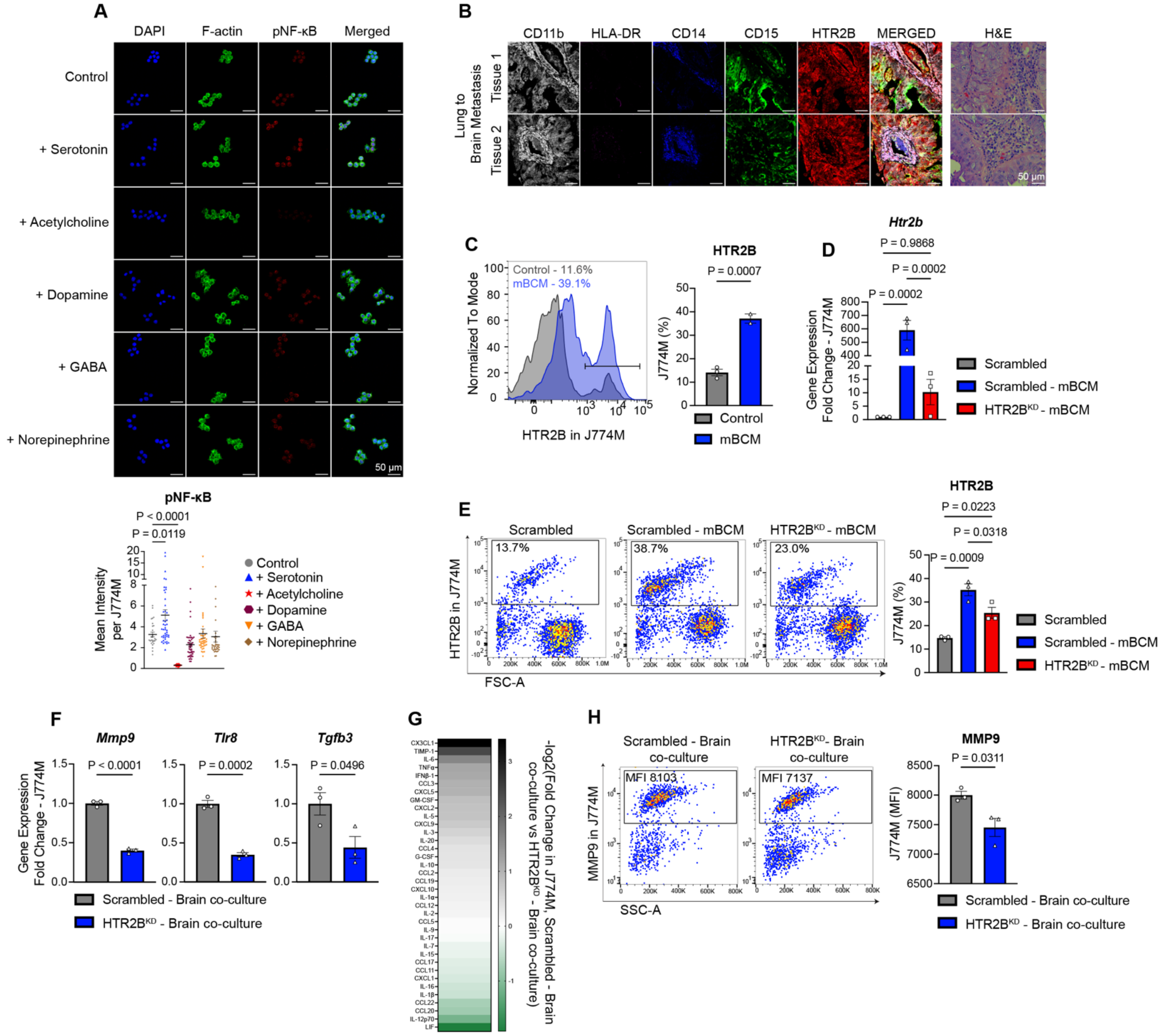


**Supplemental Figure 2: Serotonin receptor HTR2B regulates NF-κB signaling in MDSCs. (A)** IF staining and quantification of pNF-κB expression in J774M after treatment with serotonin, acetylcholine, dopamine, GABA, or norepinephrine. 3 images per group, 7-10 cells per image. Scale bar = 50 μm. **(B)** IF and H&E staining of MDSCs and HTR2B in lung-to-brain metastatic patient tissues. Scale bar = 50 μm. **(C)** Flow cytometric analysis of HTR2B expression in J774M treated with mBCM. qPCR analysis **(D)** and flow cytometric analysis **(E)** of HTR2B expression in J774M Scrambled in control media, or J774M Scrambled and HTR2B$^{KD}$ in mBCM. **(F)** qPCR analysis of NF-κB associated inflammatory markers in J774M Scrambled or HTR2B$^{KD}$ in the mMDSC-brain model. **(G)** Heatmap depicting the results of ELISA analysis comparing the levels of NF-κB associated inflammatory markers in the media of J774M Scrambled or HTR2B$^{KD}$ in the mMDSC-brain model. **(H)** Flow cytometric analysis of NF-κB associated inflammatory markers in J774M Scrambled or HTR2B$^{KD}$ in the mMDSC-brain model. Signal measured by median fluorescent intensity (MFI). Error bars represent ± SEM.

### Supplemental Figure 3

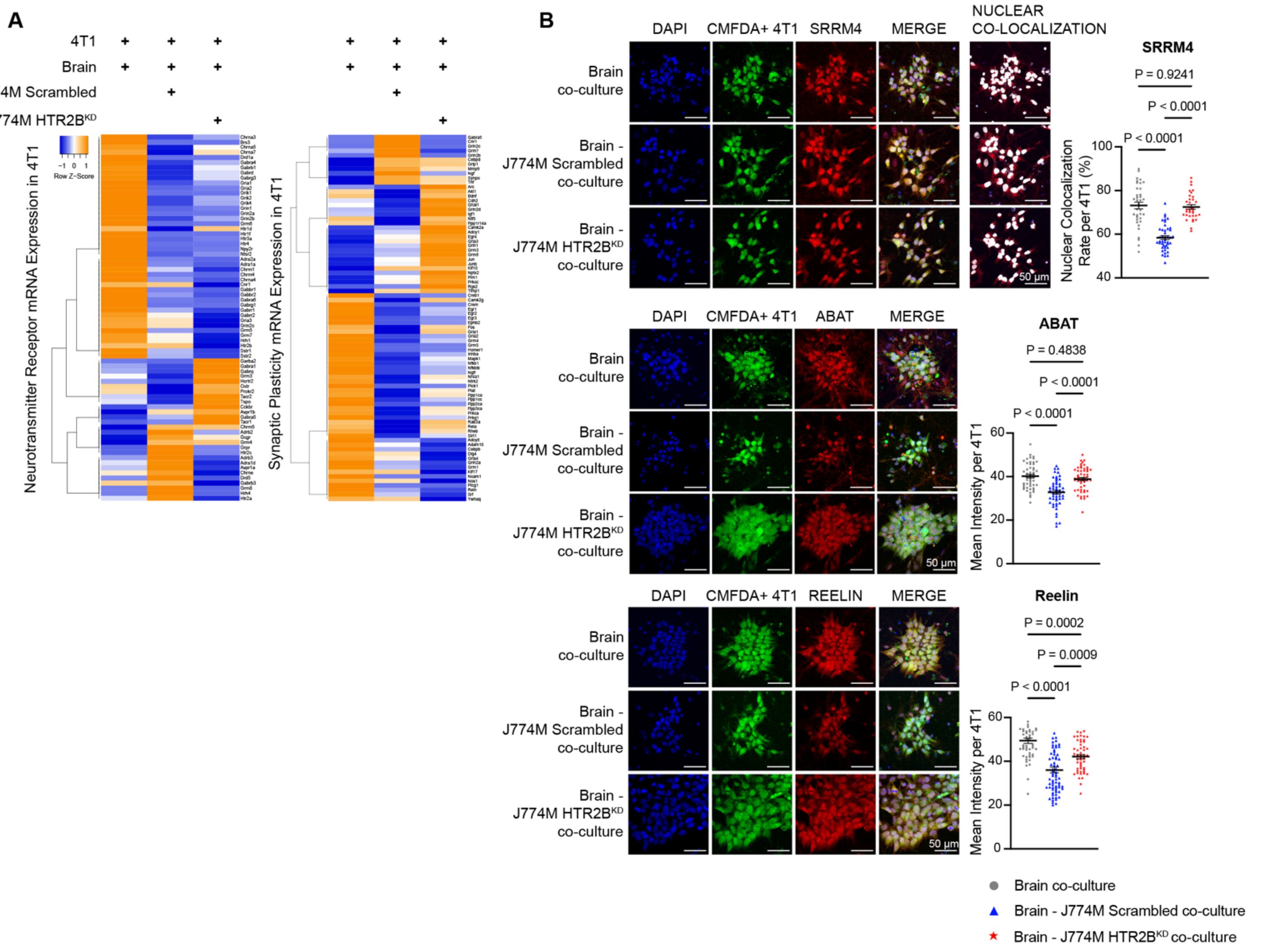


**Supplemental Figure 3: MDSC-HTR2B regulates neuronal acquisition in breast cancer cells. (A)** Clustergram depicting mRNA expression of mouse neurotransmitter receptors and synaptic mediators in 4T1 under three conditions: Brain co-culture, Brain-J774M Scrambled co-culture, Brain-J774M HTR2B$^{KD}$ co-culture. **(B)** IF staining and quantification of SRRM4, ABAT, and Reelin protein expression in 4T1 in the three conditions listed in (A). Tumor cells were stained with CMFDA dye immediately prior to co-culture seeding to distinguish from other cell populations. SRRM4 signal is measured by nuclear colocalization rate, whereas ABAT and Reelin are measured by mean intensity per cell. 4-5 images per group, 10-12 cells per image. Error bars represent ± SEM.

**Supplemental Figure 4**

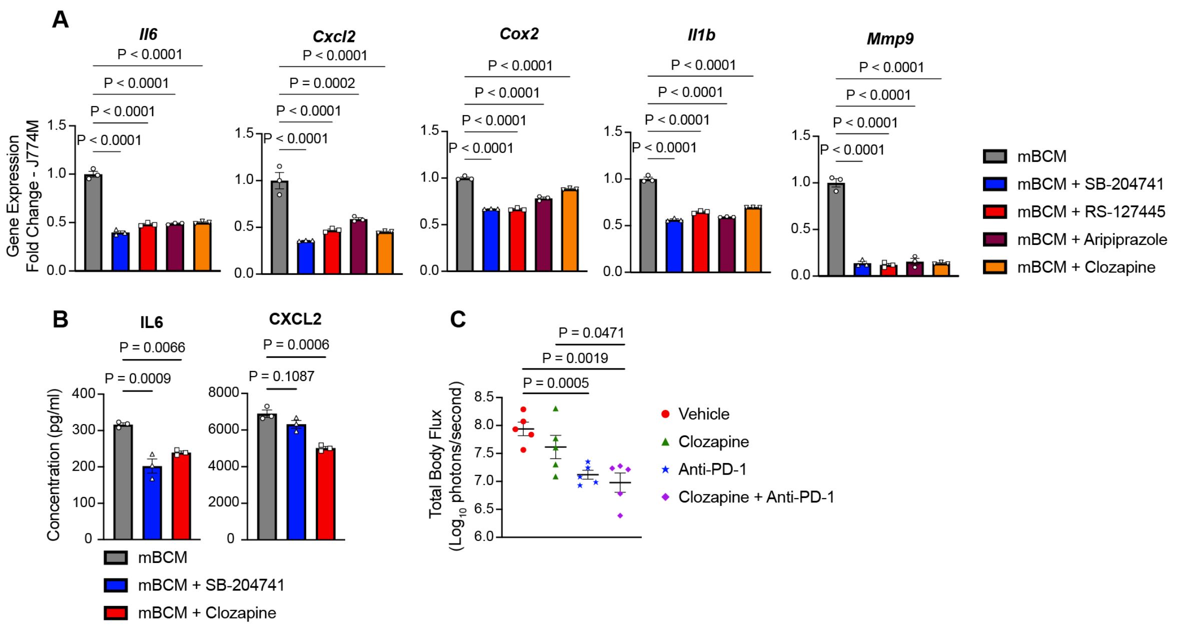


**Supplemental Figure 4: HTR2B antagonism reduces NF-κB signaling in MDSCs. (A)** qPCR analysis of NF-κB associated inflammatory markers in J774M treated with either 1 uM SB-204741, 1 uM RS-127445, 1 uM Aripiprazole, or 1 uM clozapine in mBCM. **(B)** ELISA analysis of NF-κB associated inflammatory markers in J774M treated with either 10 uM SB204741 or 10 uM clozapine in mBCM. **(C)** Bar graph represents total BLI signal at day 10 post-tumor implantation. n=5 per group. Error bars represent ± SEM.

**Supplemental Table 1**

**Primers:**

| Gene (Mouse) | Primer Sequence |
|---|---|
| *Il6* | Primer 1: 5’-AGCCAGAGTCCTTCAGAGA-3’<br>Primer 1: 5’-TCCTTAGCCACTCCTTCTGT-3’ |
| *Il1b* | Primer 1: 5'-GACCTGTTCTTTGAAGTTGACG-3’<br>Primer 2: 5'-CTCTTGTTGATGTGCTGCTG-3’ |
| *Il10* | Primer 1: 5’-ATGGCCTTGTAGACACCTTG-3’<br>Primer 2: 5’-GTCATCGATTTCTCCCCTGTG-3’ |
| *Tnf* | Primer 1: 5’-TCTTTGAGATCCATGCCGTTG-3’<br>Primer 2: 5’-AGACCCTCACACTCAGATCA-3’ |
| *Cox2* | Primer 1: 5'-CAAGACAGATCATAAGCGAGGA-3'<br>Primer 2: 5'-GCGCAGTTTATGTTGTCTGTC-3' |
| *Nos2* | Primer 1: 5'-GACTGAGCTGTTAGAGACACTT-3’<br>Primer 2: 5’-CACTTGCTCCAAATCCAAC-3’ |
| *Cxcl2* | Primer 1: 5'-CAGAAGTCATAGCCACTCCAAG-3'<br>Primer 2: 5'-CTTTCCAGGTCAGTTAGCCTT-3' |
| *Htr2b* | Primer 1: 5'-AGATTTGCTGGTTGGATTGTTTG-3’<br>Primer 2: 5'-GATGGAGGCAGTTGAAAAGAGA-3 ' |
| *Mmp9* | Primer 1: 5’-GTGGGAGGTATAGTGGGACA-3’<br>Primer 2: 5’-GACATAGACGGCATCCAGTATC-3’ |
| *Tgfb3* | Primer 1: 5’-GTGTGACATGGACAGTGGAT-3’<br>Primer 2: 5’-ACATAGGTGGCAAGAATCTGC-3’ |
| *Srrm4* | Primer 1: 5’-TCTGAAGGTCCATCCTGATCT-3’<br>Primer 2: 5’-CATCATCGTCGCCAGTATCAC-3’ |
| *Abat* | Primer 1: 5’-GGACTTCCGTCTTCATGAGTC-3’<br>Primer 2: 5’-ACCTCCACCTCTTCATACCT-3’ |
| *Reln* | Primer 1: 5’-AGCACTCTCTCCTCCTATCTG-3’<br>Primer 2: 5’-GTCGTGTCTTCTGGATCTTCTC-3’ |
| *Arg1* | Primer 1: 5'-GAATGGAAGAGTCAGTGTGGT-3'<br>Primer 2: 5'-AGTGTTGATGTCAGTGTGAGC-3' |
| *Rplpo* | Primer 1: 5’-GCACAGTGACCTCACACG-3’<br>Primer 2: 5’-AGAAACTGCTGCCTCACATC-3’ |
| *Gapdh* | Primer 1: 5’-AATGGTGAAGGTCGGTGTG-3’<br>Primer 2: 5’-GTGGAGTCATACTGGAACATGTAG-3’ |
| *Ido1* | Primer 1: 5'-GTAGAGCGTCAAGACCTGAAAG-3'<br>Primer 2: 5’-GATATATGCGGAGAACGTGGAA-3' |
| *Nfkbiz* | Primer 1: 5'-TGCAGGACCCTTACTAAGGA-3' |

| | |
|---|---|
| | Primer 2: 5'-GTTCCAGATTTGCTTCTTCCG-3’ |
| *Tlr8* | Primer 1: 5'-CGTTTTACCTTCCTTTGTCTATAGAAC-3’<br>Primer 2: 5'-TCTGGAATAGTTCGCTTTATGGA-3’ |
| *Tlr1* | Primer 1: 5'-GAGCAGAACTAGTGTTGTGAATG-3'<br>Primer 2: 5'-TCTTCAGAGCATTGCCACAT-3' |
| **Gene (Human)** | **Primer Sequence** |
| *IL6* | Primer 1: 5’-ATTCGTTCTGAAGAGGTGAGTG-3’<br>Primer 2: 5’-CCTTCCCTGCCCCAGTA-3’ |
| *IL1B* | Primer 1: 5'-CAGCCAATCTCATTGCTCAAG-3'<br>Primer 2: 5'-GAACAAGTCATCCTCATTGCC-3’ |
| *TNF* | Primer 1: 5’-TCAGCTTGAGGGTTTGCTAC-3’<br>Primer 2: 5’-TGCACTTTGGAGTGATCGG-3’ |
| *COX2* | Primer 1: 5’-TGTTTGGAGTGGGTTTCAGA-3’<br>Primer 2: 5’-GAGTGTGGGATTTGACCAGTA-3’ |
| *NOS2* | Primer 1: 5'-CACCATCCTTTGCGACA-3'<br>Primer 2: 5'-GCAGCTCAGCCTGTACT-3' |
| *CXCL2* | Primer 1: 5'-TTCACAGTGTGTGGTCAACAT-3'<br>Primer 2: 5’-TCTCGCTCTAACACAGAGGGA-3' |
| *HTR2B* | Primer 1: 5'-AGAAGAAGCTGAGTATTAC-3'<br>Primer 2: 5’-AGGCAGGACATAGAACAAGTG-3 ' |
| *RPLPO* | Primer 1: 5’-TGTCTGCTCCCACAATGAAAC-3’<br>Primer 2: 5’-TCGTCTTTAAACCCTGCGTG-3’ |

**Supplemental Table 2**

**Antibodies:**

| Primary Antibodies | | | | |
|---|---|---|---|---|
| **Antibody** | **Company** | **Catalog #** | **Host** | **Concentration used** |
| CD11b | Invitrogen | 53-0196-82 | Mouse | 1:50 |
| HLA-DR | Novus Biologicals | NB100-2707PCP | Mouse | 1:50 |
| CD14 | Novus Biologicals | NBP2-89259AF405 | Rabbit | 1:100 |
| CD15 | Abcam | ab281744 | Rabbit | 1:100 |
| CD45 | Biolegend | 103151 | Rat | 1:200 |
| CD11b | ThermoFisher | 48-0112-82 | Rat | 1:240 |
| Ly6g | Biolegend | 127605 | Rat | 1:300 |
| Ly6c | Biolegend | 128012 | Rat | 1:100 |
| pNFkB | Cell Signaling | 3033S | Rabbit | 1:500 |
| COX2 | Cell Signaling | 12282S | Rabbit | 1:200 |
| HTR2B | MyBioSource | MBS9608087 | Rabbit | 1:100 |
| CD3 | Abcam | ab16669 | Rabbit | 1:150 |
| ABAT | Novus Biologicals | NBP-2-21598 | Rabbit | 1:250 |
| SRRM4 | Biorbyt | orb2296 | Rabbit | 1:200 |
| Reelin | Genetex | GTX37552 | Rabbit | 1:200 |
| TUJ1 | R&D Systems | MAB1195 | Mouse | 1:50 |
| IDO1 | Cell Signaling | 51851S | Rabbit | 1:250 |
| NFKBIZ | Cell Signaling | 93726S | Rabbit | 1:250 |
| MMP9 | Abcam | ab38898 | Rabbit | 1:200 |
| Serotonin | Abcam | ab66047 | Goat | 1:500 |
| **Secondary Antibodies** | | | | |
| **Antibody** | **Company** | **Catalog #** | **Host** | **Concentration used** |
| Anti-Rabbit Alexa Fluor 647 | Jackson Immuno | 111-6050144 | Goat | 1:300 |
| Anti-Goat Alexa Fluor 488 | Jackson Immuno | 705-545-147 | Donkey | 1:300 |
| Anti-Mouse Alexa Fluor 647 | Jackson Immuno | 715-605-151 | Donkey | 1:300 |
